# A biological PROTAC for α-synuclein

**DOI:** 10.1101/2025.11.13.687972

**Authors:** Caitlin M. O’Shea, Gareth S.A. Wright

## Abstract

The accumulation of misfolded and aggregation-prone proteins is the hallmark of neurodegenerative diseases such as Parkinson’s disease and amyotrophic lateral sclerosis (ALS). α-Synuclein aggregation drives Parkinson’s disease pathology, and is a suitable target for selective protein clearance. Biological proteolysis targeting chimeras (bioPROTACs) aim to eliminate disease-causing intracellular proteins using host cell ubiquitination and degradation functions. Here, we describe a bioPROTAC comprising the E3 ubiquitin ligase domain of CHIP (carboxy terminus of Hsc70-interacting protein) fused to NbSyn87, a nanobody specific for α-synuclein. Co-expression with α-synuclein resulted in a significant decrease in the abundance or complete degradation of both wild-type and Parkinson’s disease-associated mutant α-synuclein (A53T, A53V, and G51D) dependent on cell type. The bioPROTAC also significantly reduced abundance of insoluble α-synuclein aggregates. In contrast, CHIP-based bioPROTACs targeting superoxide dismutase 1 (SOD1) or LIM domain only 2 (LMO2) failed to degrade their targets and in some instances, increased target abundance due to stabilising interactions with the recognition domain. These findings demonstrate key parameters for consideration during BioPROTAC design including target half-life, bioPROTAC solubility, recognition domain binding affinity, molecular chaperone activity, and interdomain linker optimisation. This work demonstrates the use of CHIP-based bioPROTACs for therapeutic degradation of α-synuclein in the synucleinopathies and provides insights for future targeted degrader development.

## Introduction

Neurodegenerative diseases (NDs) such as Parkinson’s disease (PD) and amyotrophic lateral sclerosis (ALS) are progressively debilitating conditions that place a great financial and emotional burden on healthcare systems and support networks. Misfolded insoluble aggregates and toxic soluble proteins are central hallmarks of these NDs. PD, dementia with Lewy bodies, PD with dementia and multiple-system atrophy are characterised by accumulation of misfolded α-synuclein aggregates in neuronal tissues (Dickson et al., 1999; Spillantini et al., 1998a). α-synuclein is a presynaptic neuronal protein thought to be involved in vesicle and neurotransmitter release (Nemani et al., 2010). In PD, due to both environmental and genetic factors, α-synuclein is prone to misfolding, oligomerisation and formation of insoluble fibrils each of which may a exert toxic effect on the cell (Emin et al., 2022; Mehra et al., 2019). Oxidative stress is increased by accumulation of misfolded α-synuclein, and further exacerbates cellular pathology and death. Additionally, misfolded α-synuclein can propagate to nearby cells in a prion-like manner, seeding aggregates and contributing to the widespread progressive pathology in the PD brain (Hashimoto et al., 1999; Junn and Mouradian, 2002; Prusiner et al., 2015). It is well documented that aggregated and misfolded α-synuclein is a key component of Lewy body inclusions in PD patients (Spillantini et al., 1998b).

Selectively removing misfolded or misfolding-prone protein species from cellular spaces could help to improve our understanding of the pathogenesis of diseases such as Parkinson’s disease, amyotrophic lateral sclerosis (ALS), Huntington’s disease and Alzheimer’s disease. Chemical or biological entities that can perform this function could also have therapeutic uses by addressing the shortfalls of pharmacological inhibition activity or oligomerisation. Proteolysis targeting chimeras (PROTACs) are small molecules that form ternary complexes with a target protein and endogenous E3 ligases. E3 proximity facilitates ubiquitination and proteasomal degradation of the target (Bondeson et al., 2015). Biological PROTACs (bioPROTACs) are derived from this concept but differ in that they comprise an E3 ligase ubiquitylation domain and target recognition domain fused within a single protein, meaning they are less reliant on host cell machinery (Lim et al., 2020). E3 proximity facilitates proteasomal degradation of the target protein (Bondeson et al., 2015) by bringing ubiquitylation functionality directly to the target to facilitate degradation. Recent successful applications of bioPROTACs to oncological targets, for example, KRAS (Bery et al., 2020), have fuelled application of this approach to proteins involved in neurodegenerative disease including SOD1 and α-synuclein (Carton et al., 2025; Lum et al., 2025; Vogiatzis et al., 2021, Jiang et al., 2024). However, efficient α-synuclein degradation by protein-only degraders has proven difficult to achieve (Vogiatzis et al., 2021).

Here, we describe development of a NbSyn87 nanobody-based α-synuclein-targeting bioPROTAC. NbSyn87 was used as a bioPROTAC recognition domain as it has affinity for an epitope within the highly accessible C-terminal domain of α-synuclein (Guilliams et al., 2013). The ubiquitylation domain of carboxy terminal of HSP70 interacting protein (CHIP) E3 ligase was also included as it expressed in dopaminergic neurons (Ballinger et al., 1999; Imai et al., 2002). Co-expression with α-synuclein leads to a complete or near complete α-synuclein degradation and reduction in insoluble α-synuclein species and PD-related mutants by both proteasomal and autophagic routes. Conversely, SOD1- and LMO2-specific bioPROTACs created using high affinity recognition domains showed no effect on or increased target abundance dependent on cell line despite using the same CHIP U-box ubiquitylation domain. This is despite LMO2 exhibiting similar structural disorder and lysine content in comparison with α-synuclein. We conclude that a bioPROTAC recognition domain must be soluble, bind with moderate affinity and avoid any molecular chaperone activity on the target protein.

## Results

### BioPROTAC-induced reduction of α-synuclein abundance

The anti-α-synuclein nanobody NbSyn87 recognises an epitope in the C-terminus of α-synuclein (**Fig.1A**) (Guilliams et al., 2013). Fusion of the CHIP ubiquitylation domain (residues 130-303) to the NbSyn87 nanobody aims to facilitate α-synuclein degradation (**Fig.1B)**. The predicted structure of the CHIP_130-303_-NbSyn87 bioPROTAC shows the CHIP U-box in close proximity to the substrate recognition domain (**Fig.1C**), potentially allowing ubiquitylation of α-synuclein. PROTACs and bioPROTACs must interact with their substrate to bring about degradation and α-synuclein was successfully immunoprecipitated with the myc-tagged bioPROTAC, indicating formation of a binary complex (**Supplementary Figure S1.A**).

**Figure 1.**
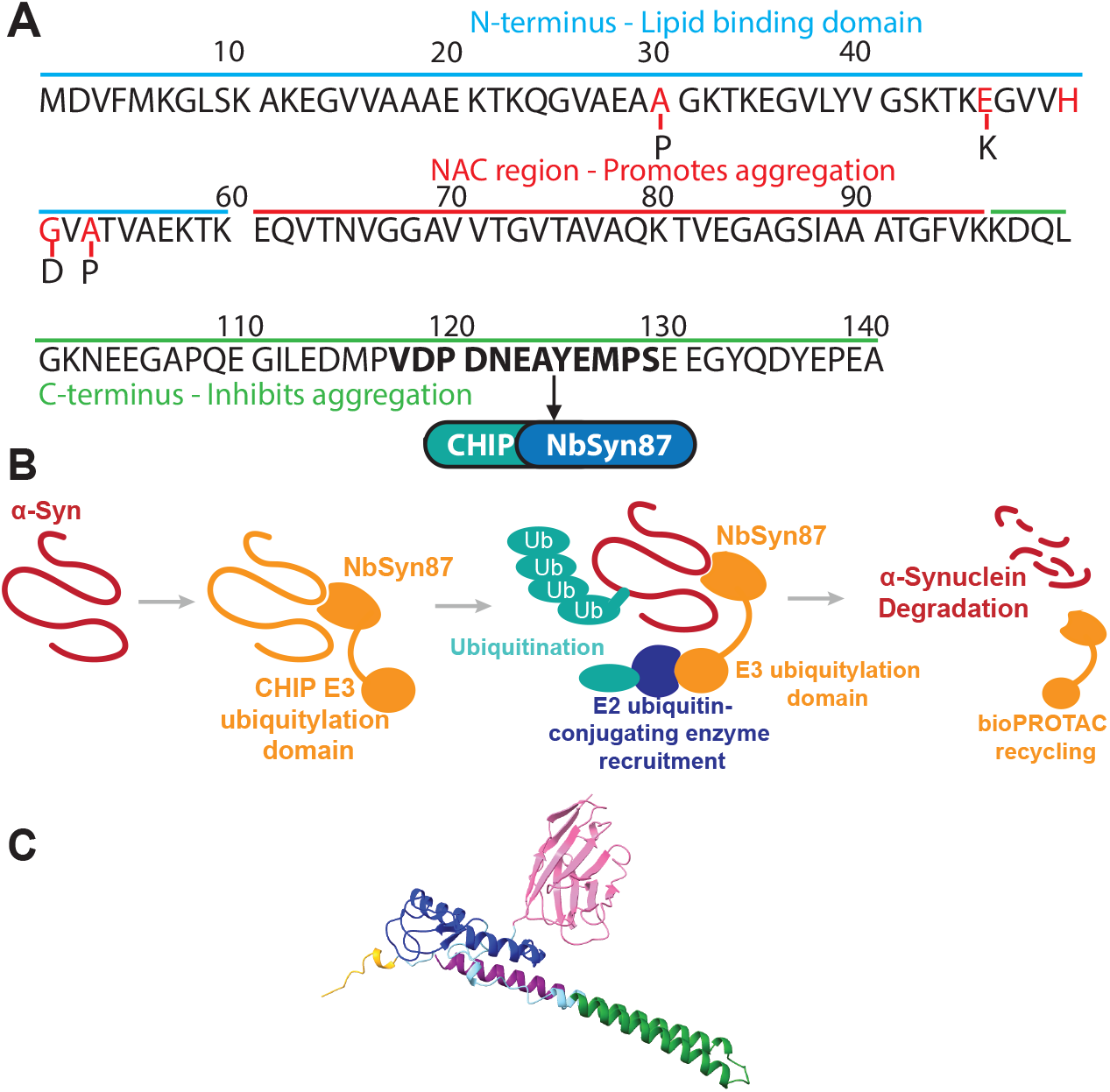
α-Synuclein sequence, bioPROTAC mechanism of action and bioPROTAC predicted structures. **A**. Schematic displaying α-synuclein sequence. Domain properties are indicated and select PD-related mutations are shown in red and region of NbSyn87 epitope recognition are shown in bold. **B**. General bioPROTAC mechanism of action. A bioPROTAC comprising NbSyn87 and CHIP_130-303_ presents bound the α-synuclein to ubiquitylation functionality encouraging degradation by host cell machinery. **C**. CHIP_130-303_-NbSyn87 bioPROTAC predicted structure.

BioPROTACs were designed with the CHIP E3 ubiquitylation domain placed at both N- or C-terminus with respect to the NbSyn87 nanobody. Placing the E3 domain at the N-terminus yielded overall better α-synuclein degradation (**Figure 2.A**) and was selected for further investigation. Overall, a 78 ± 8 % (mean ± one standard deviation, N=3) reduction in α-synuclein expression was observed (**Figure 2.B**). Non-significant degradation was observed when the NbSyn87 nanobody only was expressed with α-synuclein, in line with previous work (Butler et al., 2016). In NSC-34 cells, α-synuclein abundance was assayed as a function of bioPROTAC abundance and indicated a dose responsive relationship; as the abundance of bioPROTAC increased α-synuclein abundance decreased up to complete degradation at 1:1 coding sequence ratio (**Figure 2.C**). α-Synuclein degradation was sensitive to the cell density with incomplete degradation observed in cells at confluency at the time of harvesting (**Supplementary Figure S1.B**). This is likely due to reduced proteostatic control in non-dividing cells.

**Figure 2.**
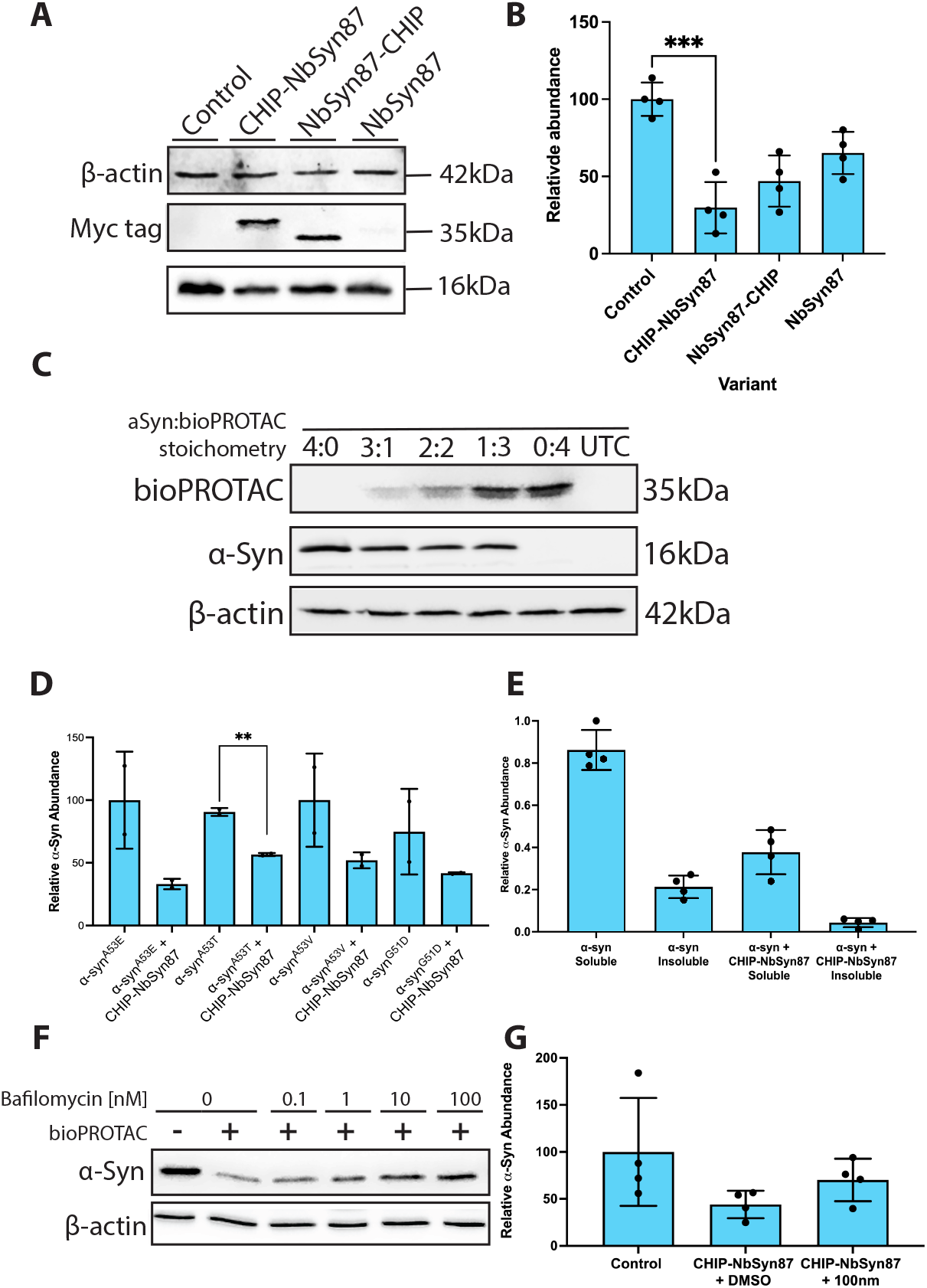
CHIP_130-303_-NbSyn87 bioPROTAC significantly reduces wild-type α-synuclein and PD-mutant abundance. **A**. Placing the E3 ligase ubiquitylation domain on the N or C terminus has different effects on α-synuclein abundance. Western blot showing reduced α-synuclein abundance when CHIP_130-303_-NbSyn87 and NbSyn87 are expressed. Co-expression of the NbSyn87 nanobody exhibited variable degradation. **B**. Mean relative abundance of α-synuclein following bioPROTAC expression. N=3, P<0.001, one-way ANOVA. **C**. α-Synuclein abundance is reduced in a dose-responsive manner, demonstrated by western blot. Plasmid DNA containing the bioPROTAC was titrated to measure any dose responsive changes to α-Synuclein abundance. β-actin was used as an internal loading control. **D**. The NbSyn87 bioPROTAC effectively reduces abundance of mutant α-synuclein. N=2. **E**. The NbSyn87 bioPROTAC significantly reduces abundance of both soluble and insoluble aggregates of α-synuclein. P<0.0001, paired t-test, N=3. **F**. α-synuclein is degraded by both autophagy and proteasomal degradation in response to NbSyn87 bioPROTAC co-expression. Western blot showing increased abundance of α-synuclein in accordance with increased concentrations of bafilomycin. **G**. Bar chart showing α-synuclein abundance when the CHIP_130-303_-NbSyn87 bioPROTAC is expressed and cells are treated with either 0 or 100 nM bafilomycin. Bar charts represent mean relative abundance values with standard deviation (error bars) and replicate data points (black).

Alanine-53 mutations in the *SNCA* gene are linked to familial PD. In particular the A53T mutation promotes misfolding of α-synuclein, leading to production of insoluble oligomers and fibrils (Polymeropoulos et al., 1997). Co-expression in HEK293T cells showed that CHIP_130-303_-NbSyn87 promotes degradation of A53T, A53V and G51D familial PD-associated mutant α-synuclein (**Fig.2D**).

### CHIP_130-303_-NbSyn87 directs α-synuclein to multiple degradation routes and inhibits its aggregation

CHIP_130-303_-NbSyn87 bioPROTAC is stably expressed in the cytoplasm (85.5% ± 5% soluble, mean ± one standard deviation, N=3) (**Supplementary Figure S1.C**). Expression of CHIP_130-303_-NbSyn87 significantly decreased the abundance of α-synuclein in insoluble aggregates, in addition to significantly reducing abundance of soluble α-synuclein (**Figure 2.E**). To determine the mechanism of α-synuclein degradation, cells transiently expressing α-synuclein and CHIP_130-303_-NbSyn87 were treated with bafilomycin, an autophagy inhibitor (Yamamoto et al., 1998). Increasing concentrations of bafilomycin gradually increased abundance of α-synuclein, suggesting less bioPROTAC-catalysed protein degradation by autophagic routes (**Figure 2.F, G**). However, abundance of α-synuclein did not fully return to control levels indicating that a portion of α-synuclein is directed to proteasomal degradation.

### BioPROTAC functionality is dependent on transient interactions with target

LMO2 is aberrantly expressed in T-cell acute lymphoblastic leukaemia and bioPROTAC-mediated degradation of this protein may impede oncogenesis. A bioPROTAC targeting LMO2 was designed using CHIP E3 ubiquitylation domain, as described above, and VH576 variable heavy domain (**Supplementary Figure S2.A**). VH576 is a known interactor and stabiliser of LMO2 (Sewell et al., 2014). Co-expression of LMO2 with the VH576 intrabody alone resulted in a significant increase in LMO2 protein abundance (**Figure 3.A**). Interestingly, no significant difference in LMO2 abundance was observed following expression of VH576 alone or CHIP_130-303_-VH576 and VH576-CHIP_130-303_, suggesting that CHIP has no effect on LMO2 stabilisation or degradation (**Figure 3.B**). This indicates that the stabilisation capabilities and high binding affinity of VH576 render it an unsuitable bioPROTAC targeting domain.

**Figure 3.**
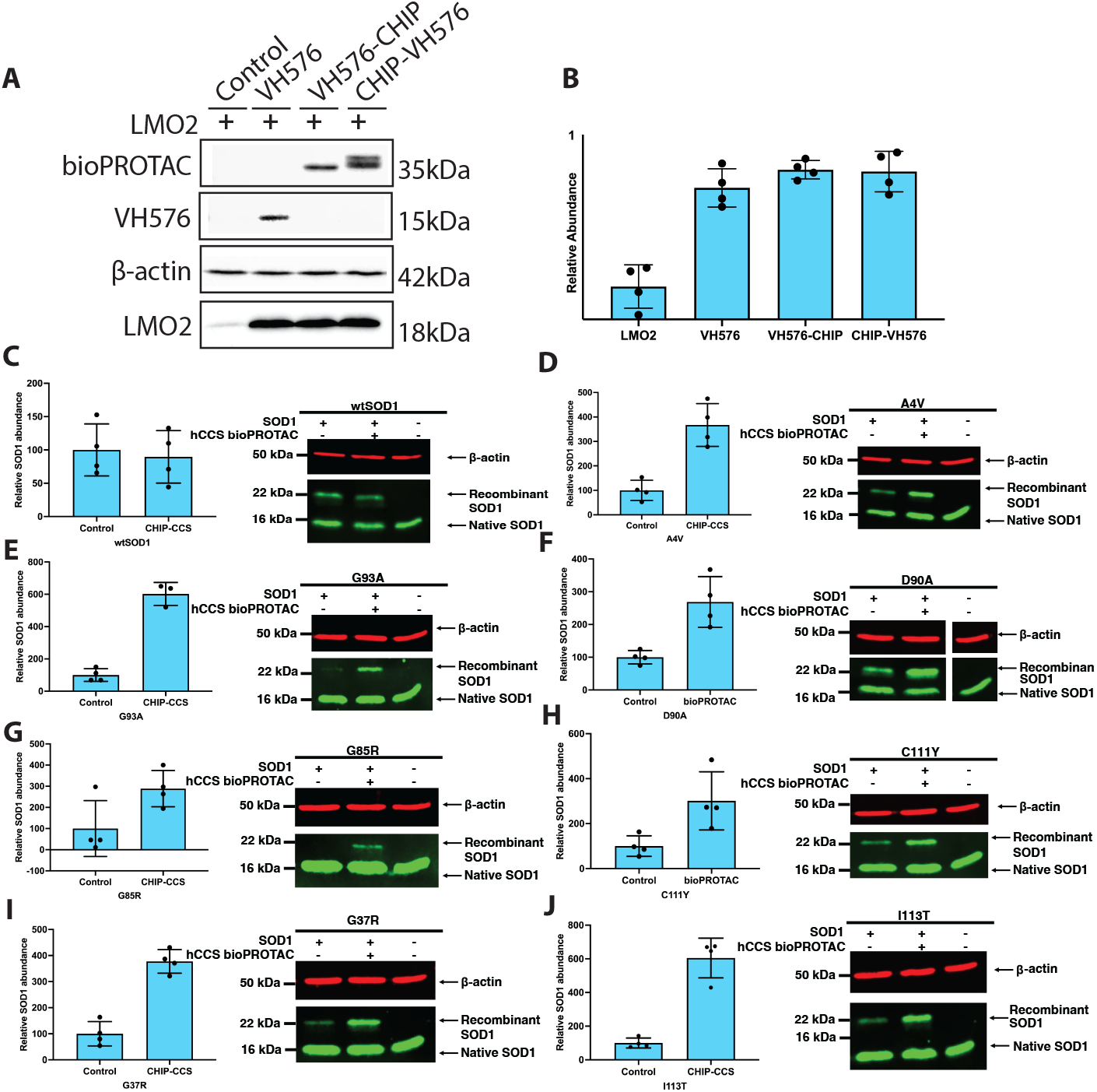
CHIP_130-303_-hCCS domain II bioPROTAC stabilises but does not degrade wildtype and ALS-mutant SOD1. **A**. No significant change to wildtype and ALS-mutant SOD1 was observed in HEK293T cells. N=4. **B**. A small but statistically insignificant decrease in SOD1 ^A4V^ and SOD1 ^G93A^ abundance was observed in HEK293S cells. N=4. HEK293T and HEK293S cells were transfected with either a wildtype or ALS-mutant SOD1 variant, or a plasmid containing a wildtype or ALS-mutant SOD1 variant and the CHIP-CCS bioPROTAC with a P2A self-cleaving peptide. and incubated for 24 hours, before sample collection and preparation for SDS-PAGE and western blotting The CHIP-hCCS bioPROTAC stabilises mutant SOD1 expression in NSC34 cells. **C**. SOD1^WT^. **D**. SOD1^A4V^. **E**. SOD1^G93A^. **F**. SOD1^D90A^. **G**. SOD1^G85R^. **H**. SOD1^C111Y^. **G**. SOD1^G85R^. **H**. SOD1^C111Y^ **I**. SOD1^G37R.^ **J**. SOD1^I113T^. Bar charts represent mean relative abundance of SOD1 following normalisation to β-actin, in which individual data points are shown, and error bars represent standard deviation. NS = non-significant. *=P<0.05, **=P<0.01, ***=P<0.005. ****=P<0.001. Paired-sample t-test, N=4. β-actin was used as an internal loading control.

A SOD1 targeting bioPROTAC was also designed using CHIP E3 ubiquitylation domain and domain II of the human copper chaperone for SOD1 (hCCS) (**Supplementary Figure S2.B**). hCCS domain II acts as a molecular chaperone for SOD1 and directly interacts with disulphide reduced wildtype and mutant or disulphide knock-out SOD1 with high affinity. Native CHIP associates with mutant SOD1 and facilitates its degradation (Choi and and Lee, 2010). hCCS domain II-CHIP_130-303_ is predominantly found in the soluble fraction indicating it is stable in the cytoplasm and available to interact with its target (**Supplementary Figure S3.A**). A second bioPROTAC was designed using CHIP E3 ubiquitylation domain and misfolded SOD1-specific nanobody Nb54, Kumar et al., (2022), but low solubility of the CHIP_130-303_-Nb54 bioPROTAC indicates poor cytoplasmic bioavailability and was predicted to be less likely to interact with and degrade SOD1 (**Supplementary Figure S3.A**). Expression of a CHIP_130-303_-CCS domain II bioPROTAC with wildtype and ALS-mutant SOD1 in HEK293T or HEK293S cells produced no significant change in SOD1 abundance (**Supplementary Figure S3.B, C**). Interestingly, a significant increase in ALS-mutant SOD1 expression was observed following expression in NSC34 cells (**Figure 3.C-J**). This increase was significant for all SOD1 mutants excluding G85R SOD1, likely due to low expression levels which are expected due to alterations in metal-binding, inability to interact with hCCS and accelerated natural protein turnover (Cao et al., 2008). No significant change in recombinant wild-type SOD1 abundance was observed again possibly due to reduced exposure to hCCS domain II due to the stability of the intrasubu nit disulphide in wild-type SOD1. To further confirm that SOD1 is stabilised rather than degraded, a selection of ALS-SOD1 mutants were co-expressed with the CHIP_130-303_-hCCS bioPROTAC in HEK293T cells and the soluble and insoluble fractions were analysed to identify changes in levels of insoluble aggregates of SOD1. Here, we observed a reduction in the abundance of insoluble SOD1 aggregates in SOD1^G93A^ and SOD1^D90A^ (**Supplementary Figure S.3D**). This demonstrates the ability of CHIP_130-303_-CCS domain II bioPROTAC to reduce levels of insoluble aggregates in HEK293T cells, which is likely a result of disulphide reduced SOD1 molecular chaperoning by CCS domain II rather than targeted degradation.

## Discussion

Effective bioPROTAC design requires careful consideration of protein target along with recognition and ubiquitylation domain selection. CHIP is highly expressed in the CNS, including in dopaminergic neurons (Ballinger et al., 1999; Imai et al., 2002). These cells must therefore express complimentary E2 conjugating enzymes. Further, CHIP has been demonstrated to ubiquitinate α-synuclein and mutant SOD1 (Urushitani et al., 2004). These features would appear to make CHIP a suitable E3 ubiquitylation domain for bioPROTACs targeting both SOD1 and α-synuclein but this is clearly not the case. Immature SOD1, LMO2 and α-synuclein show high structural disorder, accessible lysine residues, and similar polypeptide length. However, α-synuclein has a half-life of 16-48 hours (Cuervo et al., 2004; Ho et al., 2020; Mathieson et al., 2018) while LMO2 and disulfide-reduced SOD1 exhibit comparatively short half-lives of six and two hours respectively (Jonsson et al., 2006; Layer et al., 2020). α-Synuclein is efficiently degraded by our CHIP_130-303_-NbSyn87 bioPROTAC while LMO2 and SOD1 increased in abundance using an identical ubiquitylation strategy. This effect is independent of transcription or translation rates regulated by the cell given we used over-expression systems driven by the same regulatory elements. It may be that target proteins with longer half-lives are preferrable candidates for bioPROTAC degradation perhaps due to a longer window for degrader interaction, ubiquitin transfer and degradation; a concept demonstrated for small molecule PROTACs (Riching et al., 2018; Vetma et al., 2024). Furthermore, the inability to fully inhibit bioPROTAC induced α-synuclein degradation with bafilomycin indicates some non-chemotrypsin-like proteolysis. SOD1 and LMO2 may not be susceptible to these pathways. However, the most obvious difference between these examples is the impact of the recognition domain; NbSyn87 contributes to degradation while hCCS domain II and VH576 stabilise their target (Luchinat et al., 2017; Sewell et al., 2014). NbSyn87 binds to α-synuclein with high affinity, comparable to that of hCCS domain II and VH576 partner binding, suggesting that the relationship between binding affinity and bioPROTAC efficacy may be complex (Guilliams et al., 2013). NbSyn87 is not known to impact α-synuclein structure while hCCS domain II and VH576 promote disorder to order transitions by molecular chaperoning (Luchinat et al., 2017; Sewell et al., 2014). Most importantly however, NbSyn87 has a weak affinity for the proteasomal subunit Rpn10, which further drives proteasome-mediated degradation of α-synuclein, and may be a contributing factor for the efficient degradation by the CHIP_130-303_-NbSyn87 bioPROTAC (**Figure 1.B**) (Gerdes et al., 2020). Stabilisation and increased abundance of LMO2 by the variable domain VH576 and the CHIP-VH576 bioPROTAC was nonsignificant, indicating CHIP is not a facilitator of this stabilisation. From this we conclude that a bioPROTAC recognition domain must be available and therefore highly soluble, bind with moderate affinity and avoid stabilising the target protein. Substrate flexibility and target protein lysine accessibility are essential for effective ubiquitination, and substrates with few accessible lysines will be poor candidates for bioPROTAC-mediated degradation (**Supplementary Figure S4**) α-Synuclein contains several KTKGEV repeats spanning the protein, (**Supplementary Figure S4.A**), providing readily accessible sites for ubiquitination which may increase likelihood of bioPROTAC mediated degradation (Burré et al., 2014). It may also be possible that recognition domain interactions protect key lysine residues from ubiquitination by preventing access; an effect mitigated by high conformational freedom of the target. Efficient bioPROTAC mediated degradation relies on transient interactions with the target rather than persistent occupancy. High affinity stabilising interactions may prevent the bioPROTAC dissociating from a bound substrate thereby preventing bioPROTAC recycling and successive ubiquitination steps. This is key to the PROTAC concept where degrader efficiency relies not only on affinity for the target but also sub-stoichiometric catalysis (Bondeson *et al*., 2015; Mares *et al*., 2020). Similarly, endogenous E3 ligases such as CHIP rely on moderate-affinity interaction with Hsc70/Hsc90 chaperones to promote dynamic substrate ubiquitination (Murata et al 2001).

Targeted protein degradation is a promising therapeutic strategy with possible applications in neurodegenerative disease. Several small molecule PROTACs and bioPROTACs have been developed to target neurodegenerative disease targets (Cai et al., 2024). In the synucleinopathies such as PD and dementia with Lewy Bodies (DLB), cytoplasmic accumulation of misfolded and aggregated α-synuclein is central to dopaminergic neuron dysfunction and degeneration, (Lashuel et al., 2013; Spillantini et al., 1998b). Successful targeted degradation of α-synuclein by CHIP_130-303_-NbSyn87 bioPROTAC highlights a promising therapeutic avenue for PD, utilising targeted protein clearance to restore proteostasis and potentially alleviate neurodegenerative disease pathology, with the aim of delaying cell death and ultimately disease progression. It is evident that cell density impacts efficiency of observed target protein degradation (**Supplementary Figure 1B**). High cell densities may exhibit reduced proteasomal or autophagic activity and reduce protein turnover due to contact inhibition and nutrient depletion (Sabatier et al., 2025) limiting the dilution effect of cell division. Optimising cell density is therefore important when assessing degradation capabilities of PROTACs and bioPROTACs in cell models but may also present a problem in their therapeutic applications in non-dividing neuronal cell populations.

## Materials and Methods

### Cell culture

HEK293T, HEK293S and NSC34 cells were grown in DMEM supplemented with 10 % v/v foetal bovine serum (FBS), 100 units/ml penicillin and 100 µg/ml streptomycin, and 1 % v/v L-glutamine, and incubated at 37 °C with 5% CO_2_. NSC-34 cells are a fusion between primary mouse motor neurons and neuroblastoma cells offering a more relevant genetic background than HEK293T cells and may better reflect the proteostatic network found in neuronal cells (Cashman et al., 1992). Cells were split into 24-well plates at a density of 37,500-75,000 cells per well. Cells were counted using the LUNA-FL Dual Fluorescence Cell Counter (Logos Biosystems) by staining with 0.4 % trypan blue dye solution using bright field cell counting programme. For all solubility experiments and immunoprecipitation assays, cells were seeded into 6-well plates.

### DNA Transfection and Inhibitor Treatment

α-Synuclein and bioPROTAC DNA constructs were constructed from publicly available sequences and synthesised by Genscript. α-Synuclein and bioPROTAC variants in pcDNA3.4 were transfected 24 hours after cell seeding using Lipofectamine 3000 (Invitrogen) or Turbofect (ThermoFisher) according to the manufacturer’s instructions. Cells were incubated for a minimum of 16 hours and no longer than 24 hours. Bafilomycin was diluted in DMSO and added to the culture medium of cells 6 hours following transfection, and samples were incubated at 37 °C and 5 % CO_2_. DMSO was used as a vehicle control. Samples were incubated for a further 12-16 hours following treatment before sample collection.

### Protein Abundance Assay

Cells were resuspended in 100 µL PBS containing protease inhibitor cocktail (Roche) at room temperature and sonicated. 25 µL sample loading buffer (40 % v/v glycerol, 0.02 % v/v β-mercaptoethanol, 8 % w/v SDS, 0.1 % w/v bromophenol blue) was then added and samples were heated to 95 °C for five minutes. Ten microlitres of total protein lysate (10-20 µg) was loaded onto 15 % polyacrylamide gels and SDS-PAGE was run at a constant current of 80 mA for approximately 35 minutes. Proteins were transferred onto a nitrocellulose membrane using iBlot™ 2 Gel Transfer Device (ThermoFisher). Membranes were blocked with 5% w/v skimmed milk (Marvel) and incubated with primary antibodies overnight at room temperature (**Supplementary Table S1**). After washing with TBS-t (50 mM Tris-HCl, 150 mM NaCl, Tween 0.05 %) the secondary antibody was incubated for one hour with subsequent TBS-t washes before chemiluminescent detection.

### Solubility Testing

To obtain soluble and insoluble fractions, cells were resuspended in PBS containing protease inhibitor cocktail (Roche) and sonicated then centrifuged for 45 minutes at 20,000 g at 4 °C. The supernatant (400 µL) was collected, and 100 µL sample loading buffer was added (soluble fraction). The pellet was fully resuspended in 400 µL PBS and 100 µL sample buffer added (insoluble fraction) before heating at 95 ^o^C for five minutes. Samples were prepared for SDS-PAGE and analysed by western blotting as described above.

### Immunoprecipitation

All immunoprecipitation experiments were performed at room temperature, buffers were not chilled, and pH was adjusted at room temperature. After 24 hours incubation post-transfection, cells were harvested in binding buffer (25 mM Tris HCl pH 7.5, 150 mM NaCl, 1 % NP-40, 1 mM EDTA, 5 % glycerol, protease inhibitor cocktail) and incubated at room temperature for 30 minutes, pipetting up and down to resuspend cells every 10 minutes. After incubation, cells were centrifuged at 12,000 g for 10 minutes at 20 °C. Before incubation HA-Trap magnetic agarose beads (ProteinTech) with end-over-end rotation for 1 hour at room temperature. Beads were then washed three times with PBS and protease inhibitor cocktail, before elution with 4x SDS-PAGE sample buffer by heating at 95 °C for five minutes.

### Statistical analysis

Western blot densitometry quantifications were performed using ImageJ and Empiria Studio (version 3.2.0.186) and data were analysed in Prism 10 (version 10.2.3). A paired sample t-test was used to calculate significant differences between two groups. An ordinary one-way ANOVA was completed to test the significance between the means of 3 or more groups.

## Author Contributions (CRediT Classification)

**C.M.O’S**, Conceptualisation, Formal Analysis, Investigation, Visualisation, Methodology, Writing – Original Draft Preparation.

**G.S.A.W**, Conceptualisation, Formal Analysis, Funding Acquisition, Investigation, Visualisation, Methodology, Project Administration, Supervision, Writing – Original Draft Preparation.

